# Musculoskeletal design of the human shoulder: implications for neuromuscular control

**DOI:** 10.64898/2026.01.21.700644

**Authors:** Daanish M. Mulla, Dimitra Blana, Edward K. Chadwick, Peter J. Keir

## Abstract

The shoulder complex is a unique musculoskeletal structure capable of versatile motor behaviour yet requiring delicate control. The purpose of our work was to better understand the nature of musculoskeletal redundancy at the shoulder accounting for biomechanical demands and motor control strategies. Using a biomechanical model of the shoulder, we simulated a series of static exertions. Joint moment results from inverse dynamics were combined with an iterative sampling method to survey the landscape of feasible muscle activity patterns. By repeating the sampling process across different numbers of degrees of freedom at the shoulder, we demonstrate how emergent solutions are shaped by the biomechanical demands at each of the shoulder joints. Furthermore, we observed that the degree of musculoskeletal redundancy appears to be higher among the scapulohumeral muscles than the thoracohumeral and thoracoscapular muscles. Finally, we found that many of the muscle activity patterns requiring similar effort costs as the minimal effort solution have similar activation profiles, but there can be a wide range of possibilities especially at greater task intensities. Altogether, the simulations provide insight into neuromuscular control and musculoskeletal model decision-making process for the shoulder.

## Introduction

The shoulder empowers us with tremendous versatility to move our arms and exert forces across a wide workspace. As a result, we can flexibly perform a single task using several control strategies. Even in the case of static contractions, individuals can re-distribute loads within and across shoulder muscles to maintain task performance in the presence of fatigue [1, 2]. Despite exhibiting adaptable motor behaviour, our nervous system is also required to delicately balance loads across muscles to actively enforce joint stability due to the lack of passive restraints throughout the majority of the glenohumeral (GH) joint range of motion [3]. Injury to any single rotator cuff muscle can disrupt the force coupling between muscles, leading to compensatory overloads across muscles, injury progression across the rotator cuff, impaired ability to maintain GH joint stability, and functional deficits [4–8]. Thus, we are left with two pervasive, yet seemingly conflicting notions about neuromuscular control of the shoulder – one promising incredible flexibility and the second compelling tightly regulated control. To reconcile these two perspectives [9], we must ask ourselves what is the degree of neuromuscular “redundancy” (or “abundancy”) afforded to our shoulders. More directly, we want to know the extent to which muscle activity patterns are allowed to vary for a given motor task, and how these muscle load-sharing relationships are defined by the biomechanics of the shoulder complex and motor control strategies employed by our nervous system.

Grounded by a “bottom-up” physics-based perspective, neuromuscular control is shaped by the musculoskeletal design of the shoulder interacting with the environment [10]. At the muscle level, each independent muscle provides our nervous system with a control option to perform a given task. At the skeletal level, each kinematic degree of freedom (DOF) is a biomechanical constraint that dictates how our nervous system must control forces across muscles to meet the task demands (e.g., joint moment) at each DOF. Although we generally have a firm grasp of a single muscle’s functional role at a particular DOF in isolation [11, 12], the complexity (and beauty) of the shoulder is in trying to understand how the individual components integrate together for overall function. Each muscle acts upon multiple DOF crossing over potentially multiple joints. Further complicating matters, the precise force each muscle can generate and the mechanical loads that need to be satisfied at each DOF will be influenced by the state of our body and environment (i.e., posture, external forces). As such, it can be challenging to predict how muscles are required to coordinate in combination with each other without a complete model comprising the essential biomechanical components of the shoulder [3]. As a rule of thumb, the ratio between the number of muscles and DOF will determine the degree of musculoskeletal redundancy for any physically modelled system, thereby shaping the available muscle coordination strategies and robustness to muscle dysfunction [9, 13]. Based on this ratio, the upper extremity was identified as being afforded the greatest musculoskeletal redundancy among different parts of the body [13]. As acknowledged by the authors, this conclusion was based on a model without independent control of the sternoclavicular (SC) and acromioclavicular (AC) joints [14, 15], in addition to not accounting for GH joint stability requirements. Simplifying the biomechanical requirements by not capturing all relevant DOF can lead to incorrect predictions of muscle activity patterns and invalid inferences on neuromuscular control [16]. The influence of model topology on muscle coordination has been studied for lower limbs and fingers [9, 13]. How the nervous system coordinates muscles to meet the mechanical demands at each DOF is yet to be fully explored for the unique musculoskeletal design of the shoulder complex.

From a “top-down” motor control perspective, selection of specific muscle activity patterns amongst a pool of possible solutions is predominantly thought to be governed by optimization principles [10]. As applied to biomechanical models, the most common cost function is minimizing an effort-based criterion (e.g., sum of muscle activations squared). Minimal effort solutions often fail at predicting muscle co-activation strategies observed in electromyography measurements [17–20]. The differences may be due to additional neuromechanically relevant considerations, such as stiffness, that are also optimized or constraining which solutions emerge [21–25]. Further, whether our nervous system converges on optimal solutions or rather settles on “good enough” solutions is debated [10, 26–28]. There can exist a rich pool of “low effort” muscle activity patterns with marginal differences in effort costs to the minimal effort solutions [25]. The diversity of “low effort” solutions could underlie individual differences in muscle activity patterns [25]. More broadly speaking, optimization is a strong assumption that restricts the search to a single solution and blinds us to all the alternative (and perhaps more likely) muscle activity patterns that individuals may instead converge upon.

Recent advances in sampling methods now equip us with the scientific tools to evaluate the entire solution space of muscle activity patterns for high-dimensional models [26, 29–33]. Solution space investigations are a cost agnostic approach that embraces all the possible (i.e., redundant) muscle activity patterns an individual could use to complete a motor task [26, 28]. When combined with visualization techniques, they can be a powerful method for providing fundamental understanding and intuition about neuromuscular control [16, 26, 34, 35].

The purpose of our work was to investigate how biomechanical factors and motor control strategies can shape neuromuscular control of the shoulder. To do so, we used an iterative sampling method to evaluate the solution space of feasible muscle activity patterns that satisfy the mechanical constraints (i.e., joint moments) for a motor task. We investigated how the nature of this solution space is fundamentally altered by changes in biomechanical factors, namely the structure of the model (i.e., number of DOF). To evaluate how muscle activity patterns may be shaped by motor control strategies, we calculated the “cost” of each solution based on a commonly used optimality criterion. Specifically, we found the least effortful solution and sampled neighbouring solutions in the cost landscape to determine the extent to which muscle activity patterns can theoretically vary while requiring similar functional “costs” to our neuromuscular system. Revealing the family of feasible solutions through this approach allowed us to explore whether a wide range of possibilities still exist accounting for “good enough” effort considerations. Overall, we sought to understand the nature of “redundancy” afforded to our neuromuscular system at the shoulder complex accounting for biomechanical demands and motor control strategies.

## Methods

The OpenSim [36] implementation of the Delft Shoulder and Elbow Model [37] was used for all simulations. The model includes 11 rotational degrees of freedom, including 3 each at the SC, AC, and GH joints as well as 1 each at the humeroulnar (HU) and radioulnar (RU) joints. The original model comprises 137 muscle elements (i.e., musculotendon actuators). To simplify the model, the muscle set was reduced to 42 elements to represent distinct anatomical muscle sub-regions based on cadaveric investigations and prior modelling studies [12, 38, 39] (Figure 1; Table 1). The maximum isometric forces of the removed muscle elements were included in the retained elements. Each muscle element was updated to the Thelen muscle model [40].

**Figure 1:**
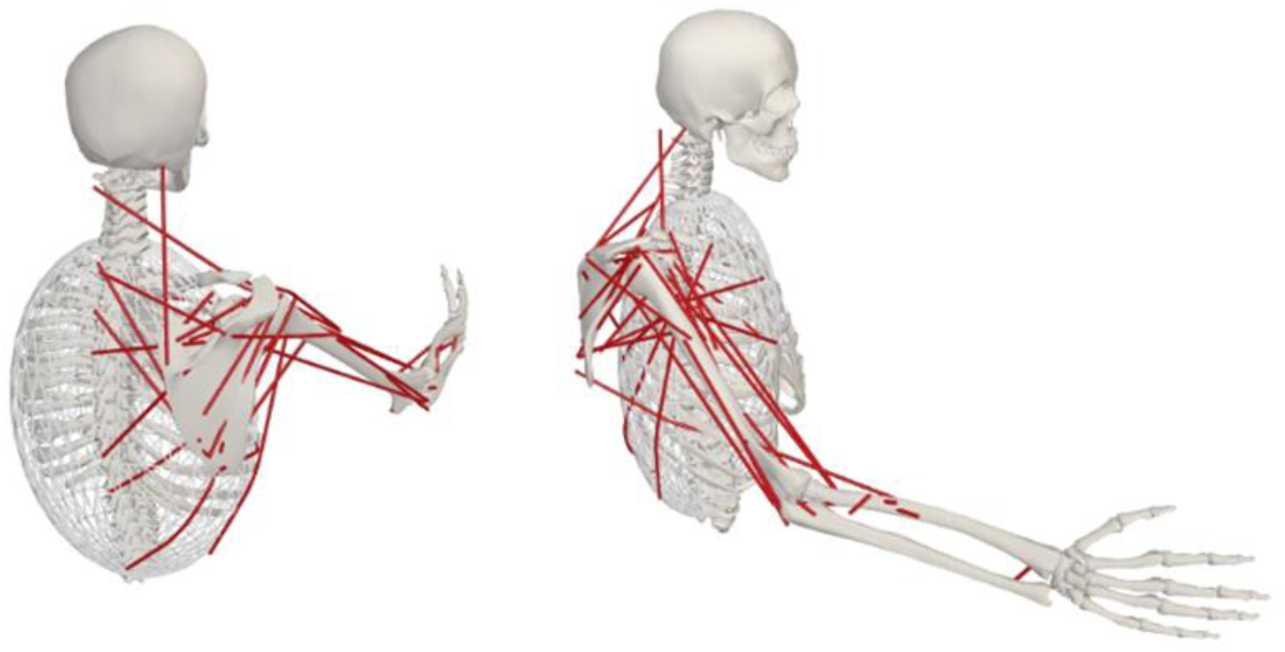
A visual representation of the Delft Shoulder and Elbow Model used in the study. The model was simplified with a reduced number of muscle elements (see Table 1). The displayed pose was used for all task conditions based on data from [41]: 60° thoracohumeral elevation in the sagittal plane (i.e., 60° shoulder flexion), elbow bent to 30° flexion, and the forearm was pronated to 45° for a medial-facing palm (i.e., thumb pointing up).

**Table 1:** The muscle elements were reduced from the Delft Shoulder and Elbow Model into distinct anatomical regions for our simplified model. For example, the *Superior Trapezius, Scapula* muscle in our simplified model had the same line of action (i.e., attachment points, wrapping set), optimal fiber length, tendon slack length, and pennation angle as the referent element *trap_scap_3*, with the maximum isometric force based on the sum of the original elements *trap_scap_1* to *trap_scap_5*.

| **Simplified Model Muscles** | **F<sub>0</sub> (N)** | **Referent (Combined) Elements** |
| --- | --- | --- |
| Superior Trapezius, Scapula | 745 | trap_scap_3 (1-5) |
| Middle Trapezius, Scapula | 336 | trap_scap_7 (6-8) |
| Inferior Trapezius, Scapula | 296 | trap_scap10 (9-11) |
| Trapezius, Clavicle | 144 | trap_clav_1 (1-2) |
| Levator Scapula | 200 | lev_scap_1 (1-2) |
| Pectoralis Minor | 307 | pect_min_2 (1-4) |
| Superior Rhomboid | 285 | rhomboid_2 (1-3) |
| Inferior Rhomboid | 149 | rhomboid_4 (4-5) |
| Inferior Serratus Anterior | 386 | serr_ant_2 (1-4) |
| Middle Serratus Anterior | 284 | serr_ant_6 (5-8) |
| Superior Serratus Anterior | 277 | serr_ant10 (9-12) |
| Posterior Deltoid | 946 | delt_scap_2 (1-3) |
| Middle Deltoid | 1856 | delt_scap10 (4-11) |
| Anterior Deltoid | 505 | delt_clav_2 (1-4) |
| Coracobrachialis | 463 | coracobr_2 (1-3) |
| Inferior Infraspinatus | 741 | infra_2 (1-3) |
| Superior Infraspinatus | 691 | infra_4 (4-6) |
| Teres Minor | 497 | ter_min_2 (1-3) |
| Teres Major | 608 | ter_maj_3 (1-4) |
| Posterior Supraspinatus | 233 | supra_1 (1-2) |
| Anterior Supraspinatus | 388 | supra_3 (3-4) |
| Superior Subscapularis | 386 | subscap_2 (1-3) |
| Middle Subscapularis | 537 | subscap_5 (4-6) |
| Inferior Subscapularis | 508 | subscap_9 (7-11) |
| Biceps Long Head | 347 | bic_l (1) |
| Biceps Short Head | 495 | bic_b (1) |
| Triceps Long Head | 1129 | tric_long_2 (1-4) |
| Superior Latissimus Dorsi | 144 | lat_dorsi_2 (1-2) |
| Middle Latissimus Dorsi | 225 | lat_dorsi_4 (3-4) |
| Inferior Latissimus Dorsi | 193 | lat_dorsi_6 (5-6) |
| Inferior Pectoralis Major | 565 | pect_maj_t_3 (1-4) |
| Middle Pectoralis Major | 331 | pect_maj_t_5 (5-6) |
| Superior Pectoralis Major | 292 | pect_maj_c_2 (1-2) |
| Triceps Medial Head | 1077 | tric_med_4 (1-5) |
| Brachialis | 1039 | brachialis_4 (1-7) |
| Brachioradialis | 164 | brachiorad_2 (1-3) |
| Pronator Teres, Humeral | 510 | pron_teres_1 (1) |
| Pronator Teres, Ulnar | 141 | pron_teres_2 (2) |
| Supinator | 838 | supinator_3 (1-5) |
| Pronator Quadratus | 470 | pron_quad_2 (1-3) |
| Triceps Lateral Head | 1144 | tric_lat_4 (1-5) |
| Anconeus | 362 | anconeus_2 (1-5) |

The tasks simulated were a static flexed posture (Figure 1) with varying forces applied at the hand. Based on available shoulder kinematics data [41], the model was set to 60° thoracohumeral elevation in the sagittal plane (i.e., 60° shoulder flexion). The elbow was bent to 30° flexion and the forearm was pronated to 45° for a medial-facing palm (i.e., thumb pointing up). Hand forces were of three magnitudes (20, 50, and 100 N) and four directions (up, down, left, and right). An unloaded task condition without applied hand forces was also simulated to investigate the effects of arm weight alone. Thus, a total of 13 total tasks (12 loaded, 1 unloaded) were simulated. Joint moments (Inverse Dynamics tool) as well as muscle moment arms and maximum isometric forces (Muscle Analysis tool) were extracted from OpenSim (version 4.3). In addition, the Static Optimization and Joint Reaction Analysis tools were used to obtain the GH joint reaction forces due to gravity and external hand load in the absence of muscle forces (i.e., model with all muscles removed).

The joint moments, muscle moment arms, and maximum isometric forces informed the mechanical static equilibrium requirements. Given a rigid linked segment system with *m* number of kinematic degrees of freedom and *n* number of actuators (i.e., musculotendon units and reserve actuators), the mechanical equilibrium equations can be expressed in linear algebra form as:

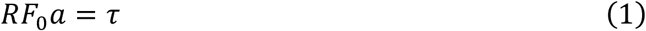

where *R* is the *m* x *n* matrix of moment arms, *F*_0_ is the *n* x *n* diagonal matrix of maximum actuator forces, *a* is the *n* x 1 vector of actuator activations, and τ is the *m* x 1 vector of joint moments [10]. Reserve actuators (maximum torque strength of 1 Nm for the GH, HU, and RU joints and 10 Nm for the SC and AC joints) were added at each degree of freedom, resulting in a total of 53 actuators (42 muscles, 11 reserves). The actuator activations are subject to the following inequality constraint from 0 (no activation) to 1 (maximum activation):

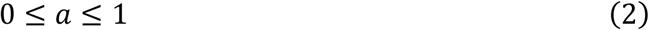

To maintain GH joint stability, the ratio of shear to compressive components of the GH joint reaction force were constrained within thresholds for preventing GH joint dislocation [42, 43]. An alternative method is to constrain the orientation of the joint reaction force within the glenoid [44. 45]; however, both approaches lead to similar conclusions on joint stability during arm elevation simulations [46], so the ratio of shear to compressive components was chosen due to ease of implementation for a linear problem The joint reaction force was calculated through the vector sum of the muscle forces crossing the GH joint (determined by the muscle lines of action from extracted muscle attachment points [12]) and the forces due to gravity and external hand loads. The GH joint reaction forces were defined in the scapular coordinate system based on International Society of Biomechanics (ISB) recommendations [47]. The coordinate system was modified such that the Z-axis (glenoid normal vector) was defined by the vector between the glenoid midpoint and root of the scapular spine (TS), as opposed to between the acromial angle and TS landmarks as per ISB conventions. We assumed that the glenoid rim was perpendicular to the modified Z-axis and thus joint compression was directed along this axis. Furthermore, the coordinate system was rotated 19° around the Z-axis to account for glenoid torsion and ensure the superior-inferior axis aligned with the glenoid poles [48].

Together, the mechanical static equilibrium equations, activation bounds, and GH joint stability requirements comprise an underdetermined linear problem where, if feasible, an infinite number of solutions (i.e., muscle activations) may exist. To represent conventional modelling approaches, we used optimization to find the solution that minimized “effort”. Effort was quantified as the sum of muscle activations normalized to the number of actuators, either as a linear (*J*_1_) or non-linear (*J*_2_) term [49]:

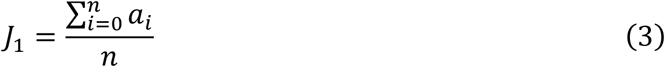

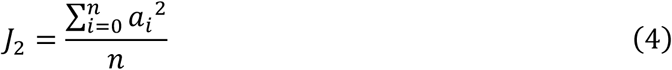

To explore the solution space, we employed two different sampling approaches using a Riemannian Hamiltonian Monte Carlo algorithm [50]. Our first sampling approach was to uniformly sample 10,000 solutions across the muscle activation landscape. We verified that 10,000 solutions were sufficient for convergence (i.e., limited further changes to the minimum and maximum muscle activations as well as correlation matrix of all sampled solutions).

Uniformly sampling solutions in the activation landscape does not (and likely will not) guarantee solutions spread across the effort landscape for such high-dimensional models. To survey across the effort landscape, our second sampling approach was to sample 100 solutions each from 100 evenly spread intervals between the minimum and maximal effort solutions (identified using optimization). This was accomplished by adding an inequality constraint to ensure solutions are only sampled within a given effort interval. As the sampling algorithm is only tractable for linear constraints, the effort intervals were based on the linear effort term (eqn. 3). In other words, we sampled a uniform distribution of solutions across the effort landscape based on the *linear effort* term. This resulted in an approximately beta distribution of solutions skewed towards a higher density of the lower range of effort values based on the *non-linear* effort term.

To investigate how the nature of musculoskeletal redundancy is shaped by limb biomechanics, we performed the sampling procedure for each simulated task on 4 models of increasing number of kinematic DOF: (i) 3 (GH joint), (ii) 5 (GH, HU, RU joints), (iii) 8 (AC, GH, HU, RU joints), and (iv) 11 (SC, AC, GH, HU, RU joints). For each model, only the joint moments from the specified DOF were incorporated into the linear problem. All results discussing the effects of the number of DOF and task demands are based on our first sampling approach (i.e., solutions sampled across the muscle activation landscape). All results discussing the effects of effort are based on our second sampling approach (i.e., solutions sampled across the effort landscape), with *non-linear* effort values reported exclusively as they are the current standard. All muscle activations are reported on a normalized scale from 0 to 100 and expressed as % maximum, which will be shortened to % for presentation purposes.

## Results

Across all simulated tasks, no feasible solutions existed for the 11 DOF model for the following exertions: 50 N (upward, rightward directions) and 100 N (all directions). If the SC joint DOF were excluded from the mechanical equilibrium constraints (i.e., 8 DOF model), feasible solutions were found for all tasks except the 100 N upward as well as 50 and 100 N rightward exertions (solutions only found with the 3 and 5 DOF models). Unless otherwise specified, we will focus on model complexities up to 8 DOF.

### Effect of number of DOF on the feasible solution space

Increasing the number of DOF decreased the range of solutions. Using the 50 N downward exertion as an example, Figure 2 depicts progressively narrower activation ranges for some muscles as the model complexity increased from 3 to 8 DOF. Uniformly distributed solutions spanning the physiological range (box and whiskers of a boxplot line up with 0, 25%, 50%, 75%, and 100% maximum) are expected for most elbow muscles (except the biceps and triceps) with the 3 DOF model and all thoracoscapular muscles for the 3 and 5 DOF models, as these muscles do not contribute to any of the mechanical equilibrium constraints for those respective models.

**Figure 2:**
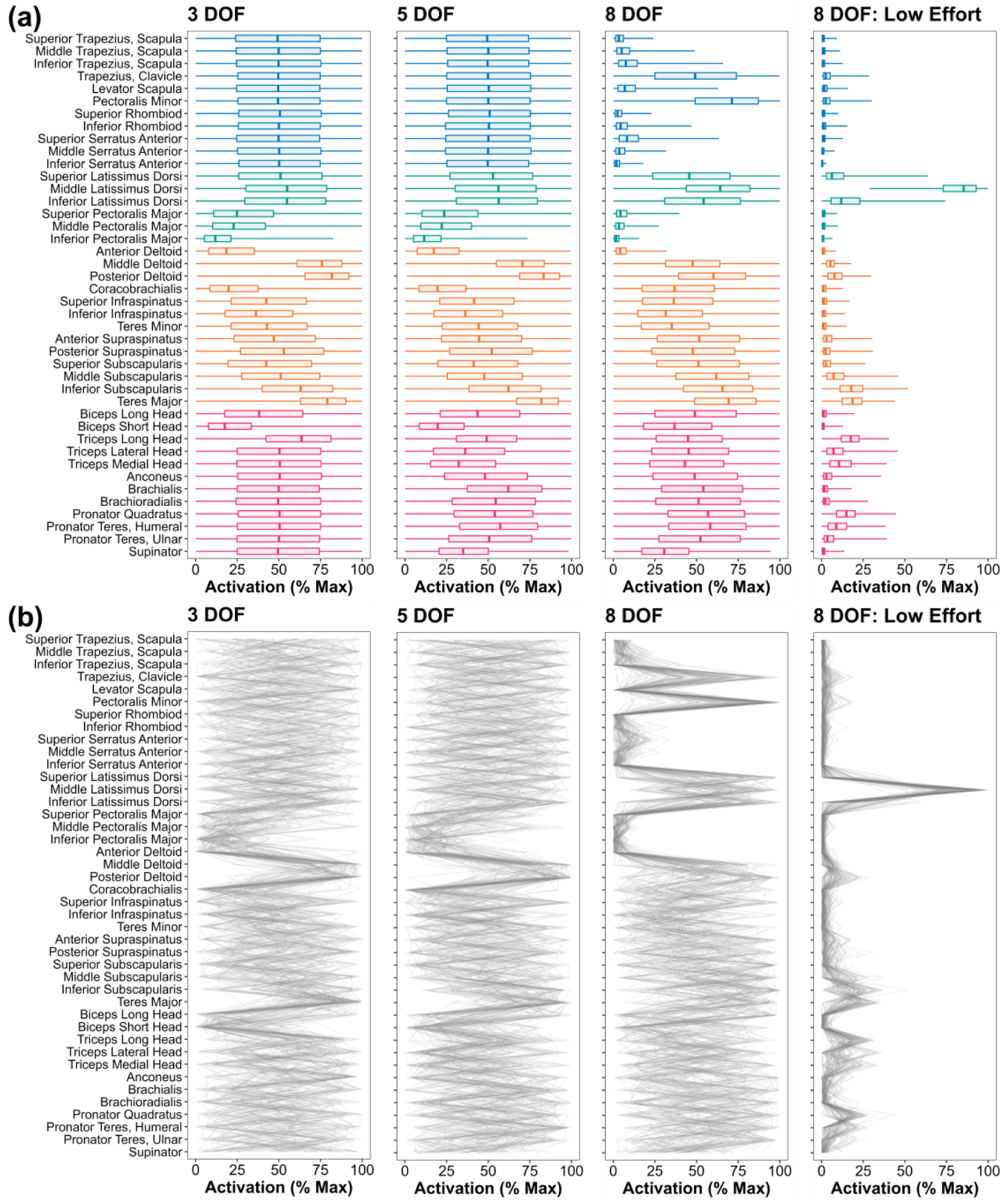
(a) Distribution of feasible solutions for a downwards 50 N exertion across models of different number of DOF and narrowing down to the 5% of solutions costing the least effort (“low effort”) based on the sum of muscle activations squared. The colours correspond to different muscle groups (blue: thoracoscapular, green: thoracohumeral, orange: scapulohumeral, pink: elbow). (b) A random sample of 100 solutions from the feasible solution space from the corresponding distributions above visualized as a flipped parallel coordinates plot. Each grey line is a single solution drawn from top to bottom (i.e., activation combination across muscles).

In the latter case, the thoracoscapular muscles only exert moments across the SC and AC joints, which are not included in the 3 and 5 DOF models. Nevertheless, even with the 8 DOF model where all muscles (except the trapezius clavicular attachment) can mechanically contribute to the task demands, the majority of muscles had solutions spanning the entire lower and upper activation bounds from 0 to 100 % maximum.

Muscle activation ranges conceal the structured families of muscle activity patterns hidden within the boundaries of the solution space. By visualizing individual activation patterns (Figure 2), we observe that despite many muscles having feasible solutions across the physiological range (0 to 100% maximum), the solution space can have internal structure, especially as model complexity increases. Of note, the thoracohumeral muscles (latissimus dorsi, pectoralis major), which are multiarticular muscles crossing the SC, AC, and GH joints, exhibit no visually apparent patterns with the 3 and 5 DOF models. Upon being required to balance AC joint moments (8 DOF model), well-defined activity patterns between the latissimus dorsi and pectoralis major muscles start to emerge (e.g., stronger, more consistent peaks and troughs visible across individual activity patterns in Figure 2). The emergent covariation between the thoracohumeral muscles with more detailed representation of the kinematic structure of the shoulder complex (i.e., adding SC and AC joints) appears across other simulated tasks (Figures S1-S3) and irrespective of lesser or greater reserve actuator strength (tested between 5-20 Nm) at the SC and AC joints (Figure S4). Similarly, the biceps and triceps, which cross the shoulder at the GH joint, display stronger visual features in the individual muscle activity patterns after inclusion of the elbow DOF for the leftward exertion (Figure S2). All these visual observations were confirmed by performing a principal component analysis on the sampled solutions. Increasing the number of DOF reduced the number of principal components required to explain greater than 90% of the variation in the solutions across muscles. The reduction in number of principal components was always equivalent to the added number of DOF, as expected with each DOF acting as a single mechanical constraint that reduces the dimensionality of the solution space by one [28].

Further examining the model with 8 DOF, we observed that the diversity of feasible activity patterns varies with muscle groups. Across all tasks, the thoracoscapular muscles encounter the narrowest activation range and display the tightest distribution of solutions. The average interquartile activation range (box of the box-and-whisker plots, i.e., middle 50% of the data) for the thoracoscapular muscles was 10.2%, compared to 24.0% (thoracohumeral), 39.6% (scapulohumeral), and 45.0% (elbow) for the other muscle groups during the 50 N downward exertion (Figure 2). This finding was consistent using other percent ranges of the data, such as examining the middle 90% of the activation range (thoracoscapular: 24.1%, thoracohumeral: 47.9%; scapulohumeral: 78.8%; elbow: 85.7%) as well as generalizable across all simulated tasks (Figure 3; Figures S1-S3). The stronger constraints on the thoracoscapular (as well as thoracohumeral) muscles are evident through visually striking features in the individual activity patterns for these groups compared to the scapulohumeral and elbow muscles, where greater stochasticity between solutions is observed (Figure 2).

**Figure 3:**
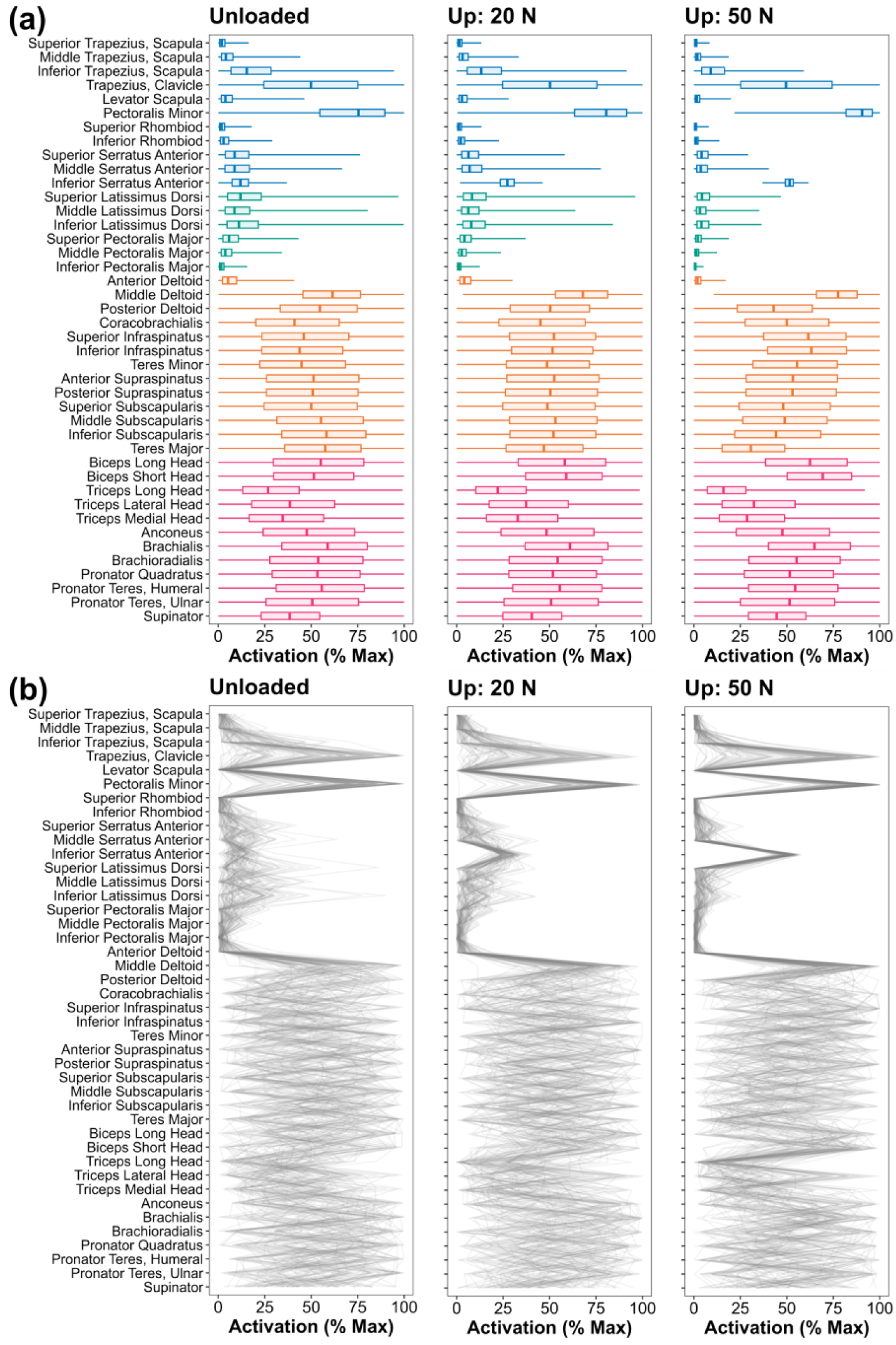
(a) Distribution of feasible solutions for tasks of increasing upwards exertion at the hand: unloaded (0 N), 20 N, and 50 N. Solutions are sampled from the 8 DOF model. The colours correspond to different muscle groups (blue: thoracoscapular, green: thoracohumeral, orange: scapulohumeral, pink: elbow). (b) A random sample of 100 solutions from the feasible solution space from the corresponding distributions above visualized as a flipped parallel coordinates plot. Each grey line is a single solution drawn from top to bottom (i.e., activation combination across muscles).

### Increasing task intensity constrains the feasible solution space

Similar to increasing the number of DOF, greater task demands will further constrain and shape the solution space. Figure 3 displays the feasible solutions progressing from an unloaded arm (0 N) to increasing hand forces applied in the upwards exertion (20, 50 N) (i.e., replicating holding increasing weights). For both the unloaded arm and 50 N upwards exertion, most muscles have feasible solutions spanning the entire physiological range (0 to 100% maximum). The average activation range across muscles decreases from 81.4% (unloaded arm) to 69.6% (50 N upwards), indicating that some muscles experience slightly narrower ranges with increasing task intensity. Moving beyond the activation ranges, the distribution of activations for many muscles skews positively (activations closer to 0) or negatively (activations closer to 100%) with increasing task intensity (see leftward or rightward shift of the boxes (interquartile range) in Figure 3 box-and-whisker plots). Closer examination of individual muscle activity patterns reveals strong rightward peaks in activations emerge for several muscles with increasing task demands (i.e., samples distributed closer to 100%). For example, only the pectoralis minor and deltoids (middle, posterior) had a median activation across all sampled solutions greater than 60% for the unloaded arm condition. Increasing the load to 50 N, additional muscles, including the infraspinatus, biceps brachii long and short heads, and brachialis, had median activations greater than 60%. Notably, these three muscles (infraspinatus, biceps brachii, and brachialis) exhibit activation ranges that nearly span from 0 to 100%. Thus, even though feasible solutions can be found with these muscles ranging from being fully relaxed to maximally active, most solutions require them to be highly activated. Although the muscles that were affected varied across task directions, we found similar findings across other tasks, with the distribution of activations being pushed to extreme ends with increased task intensity.

### Diversity of close-to-optimal (“low effort”) solutions

Searching local areas neighbouring the optimal (i.e., minimal effort) solution in the effort landscape reveals the family of “low effort” solutions that may plausibly be deemed “good enough” with respect to effort alone. Figure 4 visualizes the minimal effort solution and “low effort” solutions in both the effort and activation landscapes for a 20 N leftwards exertion.

**Figure 4:**
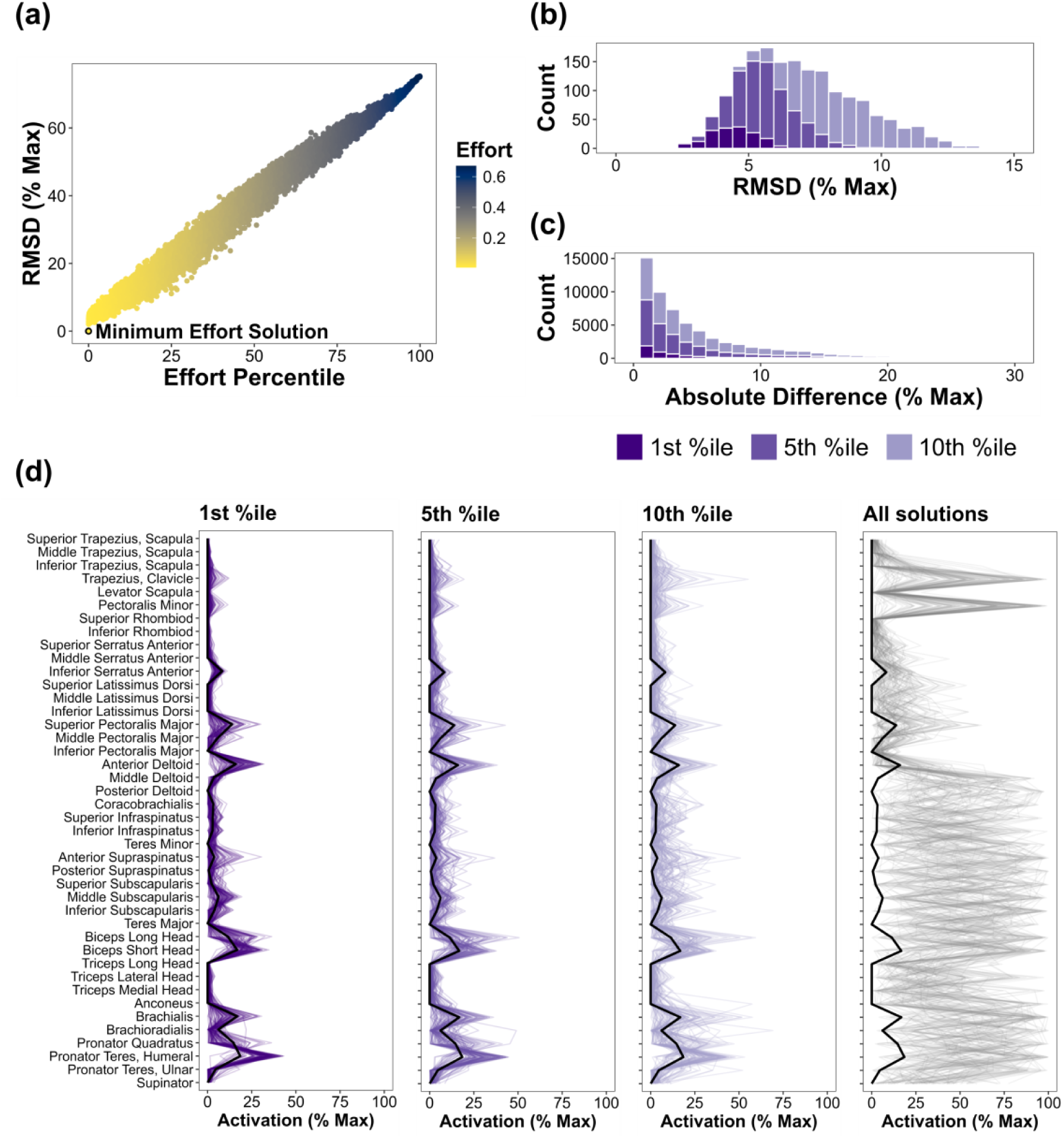
Minimum effort versus “low effort” solutions for the 20 N leftwards exertion using the 8 DOF model. (a) The RMSD in muscle activation (% maximum) between each feasible solution compared to the minimum effort solution plotted against the effort of each solution standardized to the percentile ranks of all sampled solutions in the effort landscape. The colour of the points corresponds to the effort cost of each solution (sum of muscle activations squared divided by the number of actuators). (b) Distribution of RMSD for varying “low effort” solution cut-offs (less than 1^st^, 5^th^, and 10^th^ percentile in the effort landscape). (c) Distribution of absolute difference in the activation of each muscle from the “low effort” solutions compared to the minimal effort solution. Although not visible in the histogram, we did observe rare instances where the absolute difference in muscle activation exceeded 25-50% (e.g., see brachialis in the 10^th^ percentile graph in panel d). (d) A random sample of 100 “low effort” solutions for each effort percentile cut-off (1, 5, 10%) as well as from all feasible solutions. Each line is a single solution drawn from top to bottom. The black line is the minimum effort solution.

Narrowing down to the 1% of solutions costing the least effort, the root mean square difference (RMSD) (mean ± standard deviation) in muscle activation of the sampled solutions compared to the minimal effort solution was 4.4 ± 0.9%. The RMSD increases when broadening our search to the 5% (5.4 ± 1.2) and 10% (6.9 ± 2.2) of solutions costing the least effort. For context, the mean differences in effort (sum of activations squared normalized to number of actuators) between our “low effort” solutions to the minimal effort solution were 0.005 (1% least effortful), 0.009 (5% least effortful), and 0.016 (10% least effortful), which were relatively small in magnitude compared to the range of normalized effort values across all feasible solutions (0.010 [minimal effort solution] to 0.670) (Figure 4a). We observed the diversity of “low effort” solutions increased with task intensity for the same effort percentile cut-off. Increasing the leftwards exertion to 100 N, the RMSD in activation between the “low effort” and minimal effort solutions were 9.1 ± 1.1 % (1% least effortful), 10.1 ± 1.3 % (5% least effortful), and 10.7 ± 1.5% (10% least effortful) (Figure 5). The RMSD from the 100 N leftwards exertions were the highest we observed across all simulations. Although rare, we did observe instances where the absolute difference in activation between the optimal solution and “low effort” solutions for any single muscle exceeded 25%, even when narrowing down to the 1% of solutions costing the least effort (see anterior deltoid and brachioradialis in Figure 5d).

**Figure 5:**
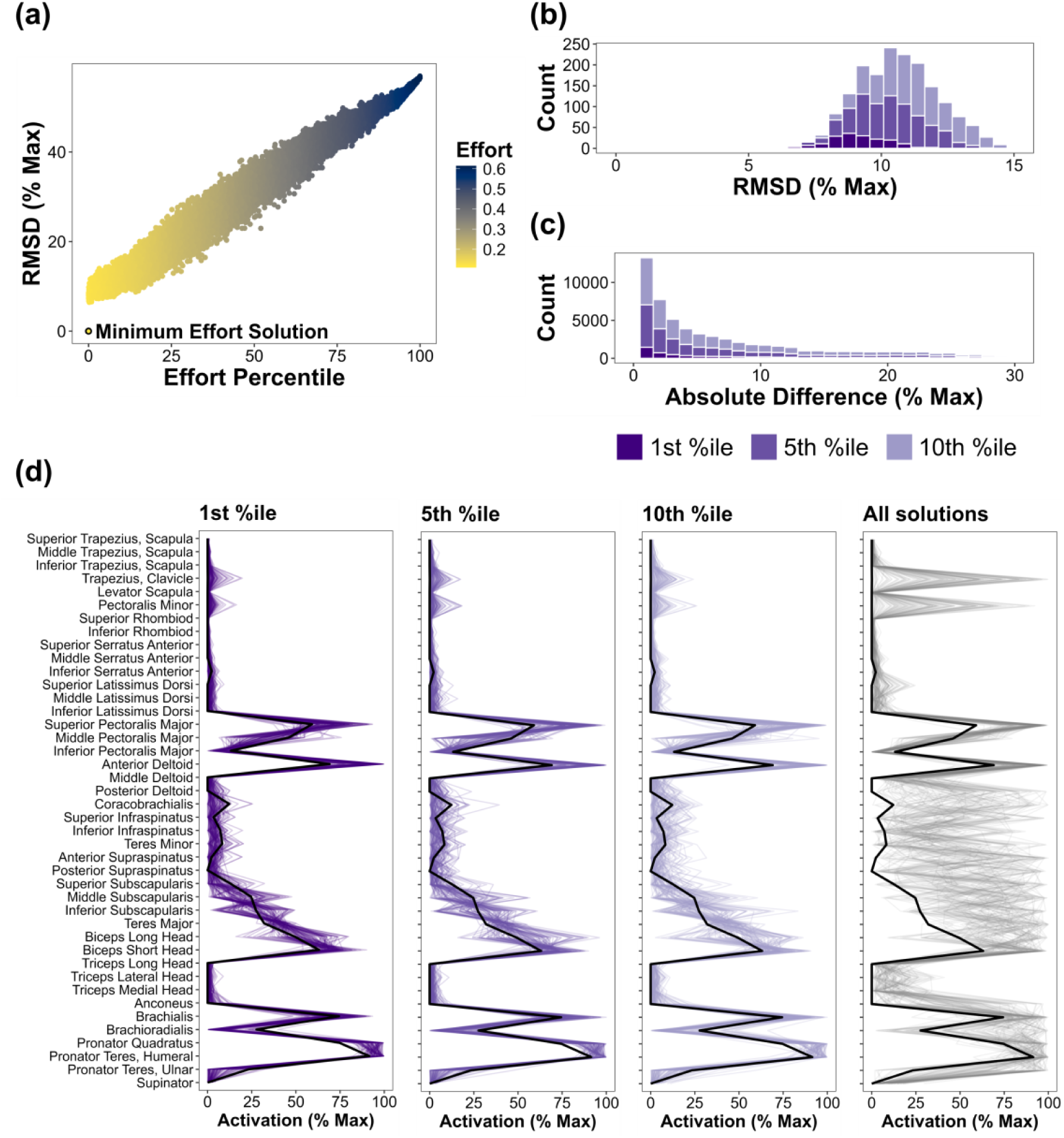
Minimum effort versus “low effort” solutions for the 100 N leftwards exertion using the 8 DOF model. (a) The RMSD in muscle activation (% maximum) between each feasible solution compared to the minimum effort solution plotted against the effort of each solution standardized to the percentile ranks of all sampled solutions in the effort landscape. The colour of the points corresponds to the effort cost of each solution (sum of muscle activations squared divided by the number of actuators). (b) Distribution of RMSD for varying “low effort” solution cut-offs (less than 1^st^, 5^th^, and 10^th^ percentile in the effort landscape). (c) Distribution of absolute difference in the activation of each muscle from the “low effort” solutions compared to the minimal effort solution. Although not visible in the histogram, we did observe rare instances where the absolute difference in muscle activation exceeded 50% (e.g., see biceps brachii long head in the 10^th^ percentile graph in panel d). (d) A random sample of 100 “low effort” solutions for each effort percentile cut-off (1, 5, 10%) as well as from all feasible solutions. Each line is a single solution drawn from top to bottom. The black line is the minimum effort solution.

## Discussion

In this study, we investigated how the musculoskeletal design of the shoulder and motor control strategies driven by effort considerations shape neuromuscular control of the shoulder. Using a combination of biomechanical modelling, high-dimensional sampling methods, and visualization techniques [25, 28, 35, 50], we provide an extensive description of the landscape of muscle activity patterns for static exertions. Our results demonstrate how muscle activity patterns are successively shaped by each joint of the shoulder complex, the degree of redundancy afforded to different muscle groups, and reveal the family of “low effort” solutions that individuals may converge upon if guided by “good enough” effort criterion. Overall, our work has far-reaching insight from neuromuscular control to musculoskeletal model decision-making process that will be discussed.

Neuromuscular control of the shoulder is shaped by the several DOF comprising the shoulder complex. By adding the moment constraints from each joint in a sequential manner, we found progressively smaller muscle activation ranges and altered load-sharing patterns. Given the multiarticular nature of many shoulder muscles, changes to one muscle do not occur in isolation and require compensatory responses to balance all joint moments [3]. As a clear example, after adding the AC joint to the mechanical constraints, the distribution of activations for the latissimus dorsi shifted positively towards greater magnitudes while requiring the anterior deltoid to be minimally active for the simulated 50 N downward exertion (Figure 2). This observation matches empirical data [51] and demonstrates an important principle. Muscle activity patterns can emerge simply from meeting the biomechanical demands of the task before even considering any “top down” motor control strategies (e.g., optimization, synergies) [26]. In the aforementioned example, the activation range for the latissimus dorsi did not change with the addition of the AC joint (0 to 100% maximum), highlighting the limitations of only examining the lower and upper bounds of feasible solutions [52, 53]. Capturing the landscape of all solutions allows us to tease apart how our nervous system is required to coordinate muscles as determined by biomechanical constraints versus exploiting families of solutions within what is feasible as governed by motor control strategies.

In addition to gaining insight into the influence of musculoskeletal design on neuromuscular control, we can simultaneously inform the model decision making process. All models require assumptions, with simplifications often made to the shoulder complex, such as not explicitly balancing moments at the SC and AC joints [14, 15, 54]. As we observed, without balancing AC joint moments in the 3 and 5 DOF models compared to the 8 DOF model, the functional demands of the thoracohumeral muscles can be fundamentally altered which may have downstream effects on the scapulohumeral muscles. If the scientific question is one of understanding muscle coordination, care must be taken that the biomechanical requirements of muscles are not removed, leading to erroneous conclusions of neuromuscular control [55].

Successively increasing the number of DOF severely limits the feasible muscle activity patterns to the point that solutions were not found after including the SC joint for half the simulated tasks. In other words, accounting for all DOF spanning from the SC joint to the elbow, muscles (by themselves) are not sufficient to generate the torques to balance out all the joint moments. At immediate glance, the fragility of the shoulder model to find feasible solutions supports the argument that our musculoskeletal system is not as “redundant” as commonly believed. Rather, we may barely have the number of muscles and DOF to function [28]. However, it is important to acknowledge that the simulations were purely muscle-driven and do not include the passive moment contributions from ligaments nor the scapulothoracic reaction forces from the closed-chain mechanism of the shoulder complex [3, 45, 56]. Our results reinforce the notion that the scapulothoracic constraints are essential for control of the shoulder [45]. Constraints from ligaments and the scapulothoracic gliding plane will reduce the dimensionality between the SC and AC joints down from 6 independent DOF [45, 56, 57]. As such, the musculoskeletal redundancy afforded at the shoulder is likely closer to the 8 DOF model simulations presented throughout our results than the over-constrained 11 DOF model.

Evidenced by the wider activation range and less well-defined activation patterns (Figures 2 and 3), there appears to be greater redundancy across the scapulohumeral muscles compared to the thoracohumeral and thoracoscapular muscles. It is possible that the missing contributions from passive tissues and scapulothoracic reaction forces over-constrain the thoracohumeral and thoracoscapular muscles with the 8 DOF model. To represent these missing components, we included reserve actuators (1 Nm at the GH, HU, and RU joints; 10 Nm at the SC and AC joints) as a “catch-all” extra moment contributor should muscles not be sufficient for satisfying the net joint moments. The strength of these reserve actuators is typical of what is required for satisfying the inverse dynamics moments across upper extremity models that incorporate SC and AC joints [45, 58]. We found no meaningful difference in the muscle activation range or distribution of solutions when repeating our simulations with decreased or increased SC and AC joint reserve actuator strength from 5 Nm up to 20 Nm. On a similar note, the GH joint is predominantly muscle-driven and the activation ranges may be inflated due to excessive reserve actuator strength. However, decreasing the actuator strength at the GH joint from 1 Nm to 0.1 Nm likewise did not substantially alter the muscle activation ranges or distribution of solutions. Thus, we are reassured that the observed difference in musculoskeletal redundancy between muscle groups is not solely a by-product of missing non-muscle moment contributors and choice in reserve actuator strength. It is not completely clear to us why there may be greater redundancy across the scapulohumeral muscles, which appeared to be particularly prominent among the rotator cuff muscles. As the rotator cuff muscles are the primary stabilizers of the GH joint and prone to injury [12, 59], perhaps the greater musculoskeletal redundancy afforded is to protect against overloading. Modelling and cadaveric studies demonstrate GH joint stability can be maintained with tears affecting a single rotator cuff muscle, consistent with epidemiological evidence of majority tears being asymptomatic, but this may come at the expense of greater loads amongst the rest of the rotator cuff group [6, 60, 61].

Consequently, tear propagation to a second rotator cuff muscle leads to GH joint instability [6, 61]. Furthermore, our model did not consider endpoint stiffness, which can be an important factor in shaping muscle activity patterns for maintaining task performance [22, 23, 33, 62]. It may be that the scapulohumeral and elbow muscles play a larger role in regulating endpoint stiffness, and therefore, the seemingly greater redundancy among these muscles diminishes once we account for stiffness control.

After accounting for biomechanical constraints, the exploitation of “low effort” control strategies further confines the feasible solutions available to our nervous system. The diversity of “low effort” solutions will depend on what we deem as “good enough”. The more restrictive the criteria (i.e., lower effort percentile cut-off), the fewer the possibilities. We observed a linear relationship between how “close” all feasible solutions were to the minimal effort solution in the effort landscape to their distance (i.e., RMSD) in the activation landscape (Figures 4a, 5a).

Similar to the observations by Sohn & Ting (2018) [25], we noted activation differences for a single muscle could differ up to and beyond 25% (Figures 4c, 5c) even though the effort differences appear insubstantial. Building upon their work [25], we also quantified the distribution of the “low effort” solutions in addition to examining the activation range. In doing so, we found that most solutions tend to be rather “close” in muscle activation space, particularly for less intensive tasks (RMSD < 10% maximum to the minimal effort solution and similar activation profiles) (Figures 4, 5). However, differences in muscle activations between “low effort” solutions grew with more intensive tasks. This finding is in line with recent hand modelling simulations, where the 5 solutions generating the strongest pinch force exhibited muscle activation differences an order of magnitude larger than pinch force differences [20]. The diverse pool of close-to-optimal solutions may be one of the drivers for between-individual variations in muscle activity patterns [25].

The exciting next step is to map emergent solutions onto the theoretical activation and effort landscapes. By identifying how individuals navigate and occupy the landscape of feasible solutions, we can generate increasingly focused null models to pinpoint the biomechanical factors and motor control strategies that shape muscle activity patterns [63]. For instance, if individuals are converging on “good enough” motor control strategies based on an effort criterion alone, we should expect to see similarities between emergent solutions and the family of feasible solutions with comparable effort costs. However, if emergent solutions occupy a narrower range than equivalently effortful solutions, this could indicate that some of these “good enough” solutions with respect to effort alone are more attractive than others. Perhaps other neuromechanical factors, such as stiffness, may then be shaping muscle coordination strategies. By considering the entire solution space of muscle activity patterns, the stage is now set to tackle these questions empirically [26] and reconcile the long-standing discrepancies between model-predicted and experimentally measured muscle activity patterns that continues to plague the most advanced upper extremity models available today.

In addition to the limitations of a purely muscle-driven shoulder model, some considerations need to be made when interpreting the study results. As with any biomechanical model, the simulations are sensitive to model parameters [55, 64, 65]. Variations in muscle model parameters will affect muscle activations predictions. Although the specific activation magnitude of minimal effort and “low effort” solutions may vary, the overarching outcomes of the study are generalizable across parameter sets (e.g., altered load-sharing relationships between muscles with number of DOF). The model assumes muscle elements (including different elements of the same muscle) behave independently. The extent to which neuromechanical factors constrain the activity patterns between and within muscles is heavily debated [10].

Modelling muscle dependencies could be implemented by enforcing activation relationships into the linear problem [66] but needs to be done with caution as these constraints may not necessarily be mechanistic [28, 67, 68]. Finally, only static exertions were simulated and results may not necessarily generalize to dynamic conditions. Recent work demonstrates that the added temporality of activation-contraction dynamics will further constrain the feasible solutions [29].

In conclusion, sampling from the entire landscape of feasible solutions revealed important insight into the neuromuscular control of the shoulder complex. The degree of musculoskeletal redundancy at the shoulder varies across different muscle groups and is sequentially constrained by each DOF in addition to effort-driven motor control strategies. Accounting for biomechanical factors and effort-driven considerations, there remain a large pool of feasible solutions. Moving forward, locating real-world muscle activity patterns onto the solution space could help in determining the neuromechanical factors shaping emergent solutions from the broader landscape of all possibilities.

