## Supplementary File for "Musculoskeletal design of the human shoulder: implications for neuromuscular control"

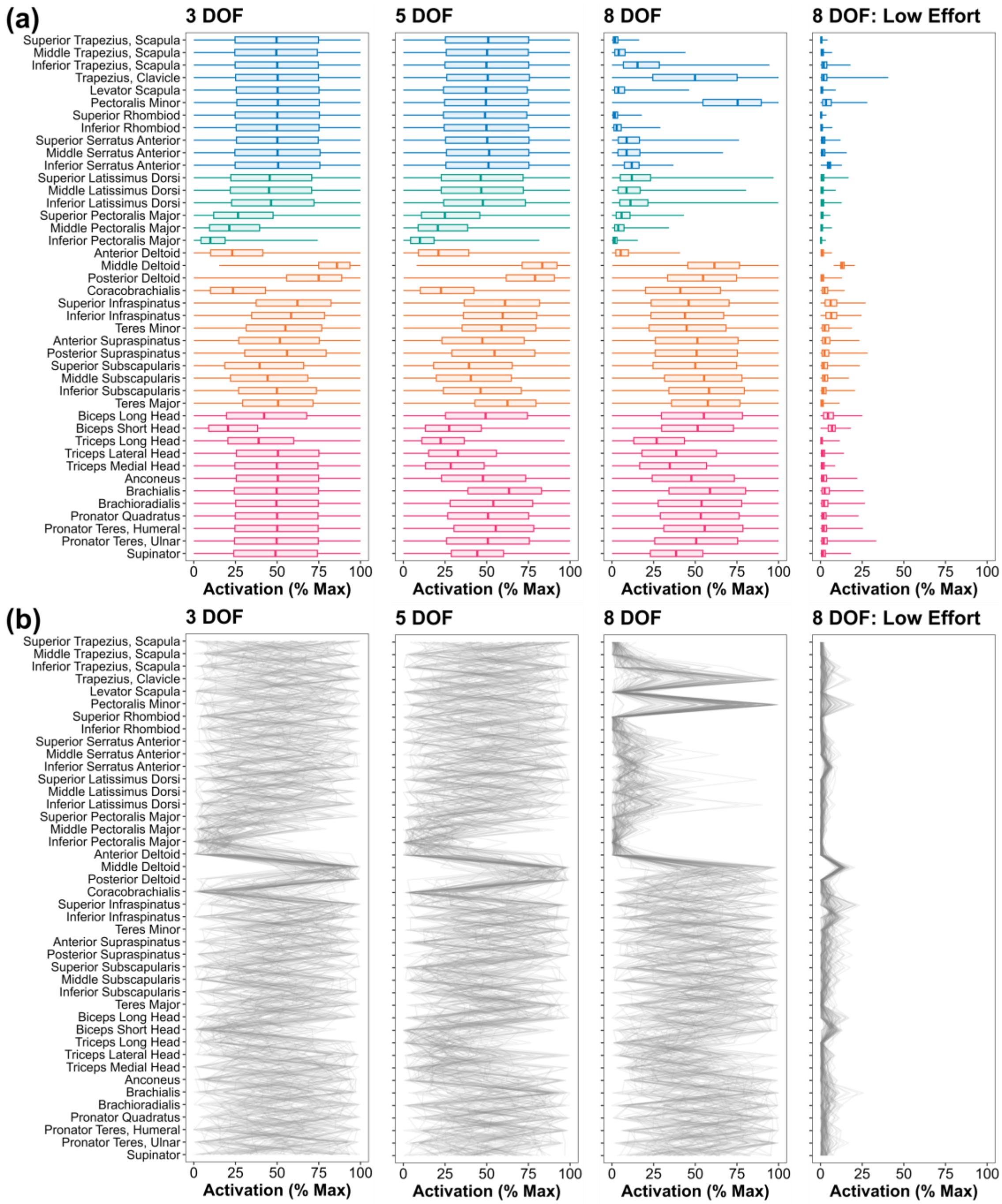


**Figure S1:** (a) Distribution of feasible solutions for an unloaded arm (i.e., holding pose) across models of different number of DOF and narrowing down to the 5% of solutions costing the least effort ("low effort") based on the sum of muscle activations squared. The colours correspond to different muscle groups (blue: thoracoscapular, green: thoracohumeral, orange: scapulohumeral, pink: elbow). (b) A random sample of 100 solutions from the feasible solution space from the corresponding distributions above visualized as a flipped parallel coordinates plot. Each grey line is a single solution drawn from top to bottom (i.e., activation combination across muscles).

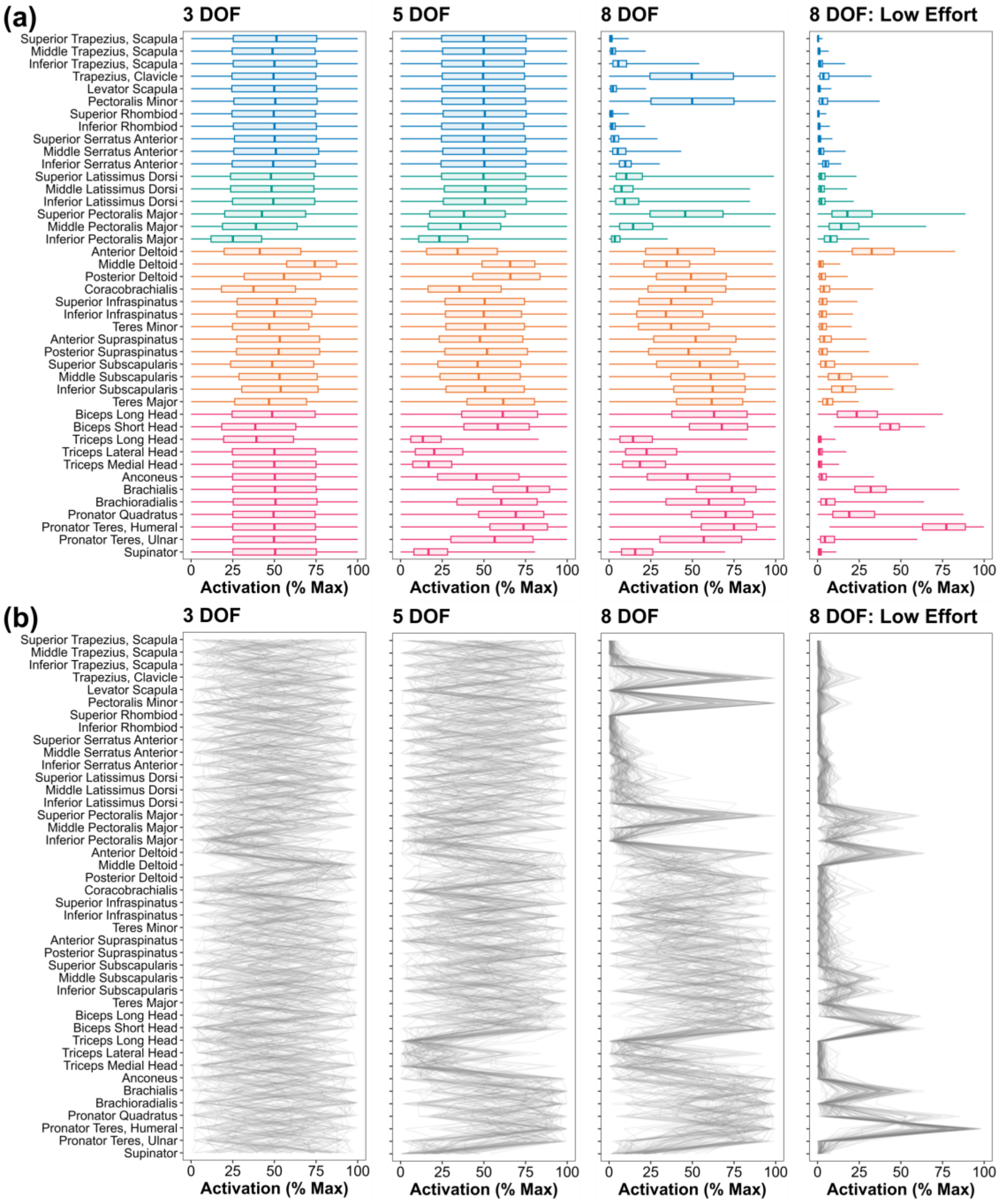


**Figure S2:** (a) Distribution of feasible solutions for a 50 N leftwards exertion across models of different number of DOF and narrowing down to the 5% of solutions costing the least effort ("low effort") based on the sum of muscle activations squared. The colours correspond to different muscle groups (blue: thoracoscapular, green: thoracohumeral, orange: scapulohumeral, pink: elbow). (b) A random sample of 100 solutions from the feasible solution space from the corresponding distributions above visualized as a flipped parallel coordinates plot. Each grey line is a single solution drawn from top to bottom (i.e., activation combination across muscles).

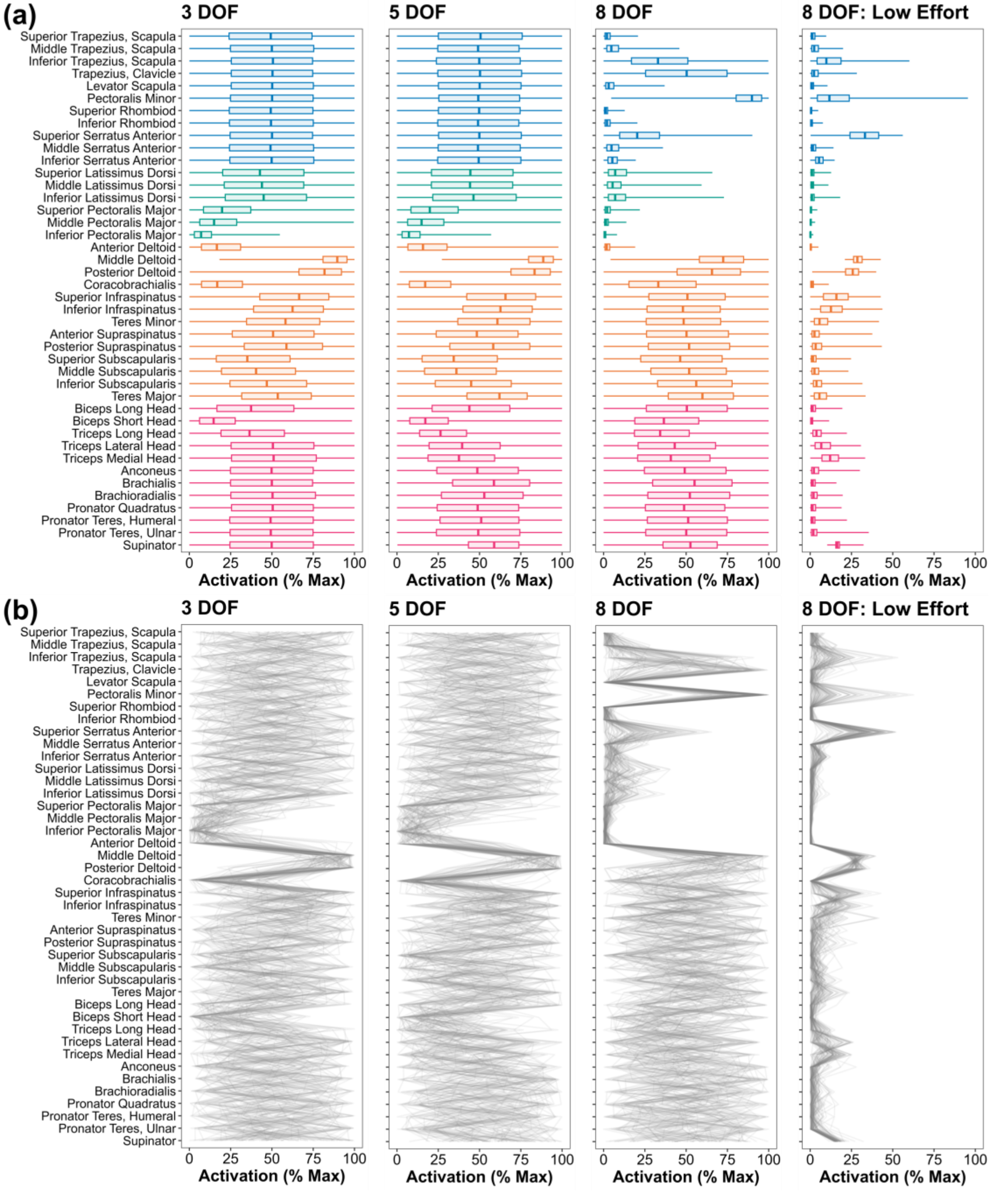


**Figure S3:** (a) Distribution of feasible solutions for a 20 N rightwards exertion across models of different number of DOF and narrowing down to the 5% of solutions costing the least effort ("low effort") based on the sum of muscle activations squared. The colours correspond to different muscle groups (blue: thoracoscapular, green: thoracohumeral, orange: scapulohumeral, pink: elbow). (b) A random sample of 100 solutions from the feasible solution space from the corresponding distributions above visualized as a flipped parallel coordinates plot. Each grey line is a single solution drawn from top to bottom (i.e., activation combination across muscles).

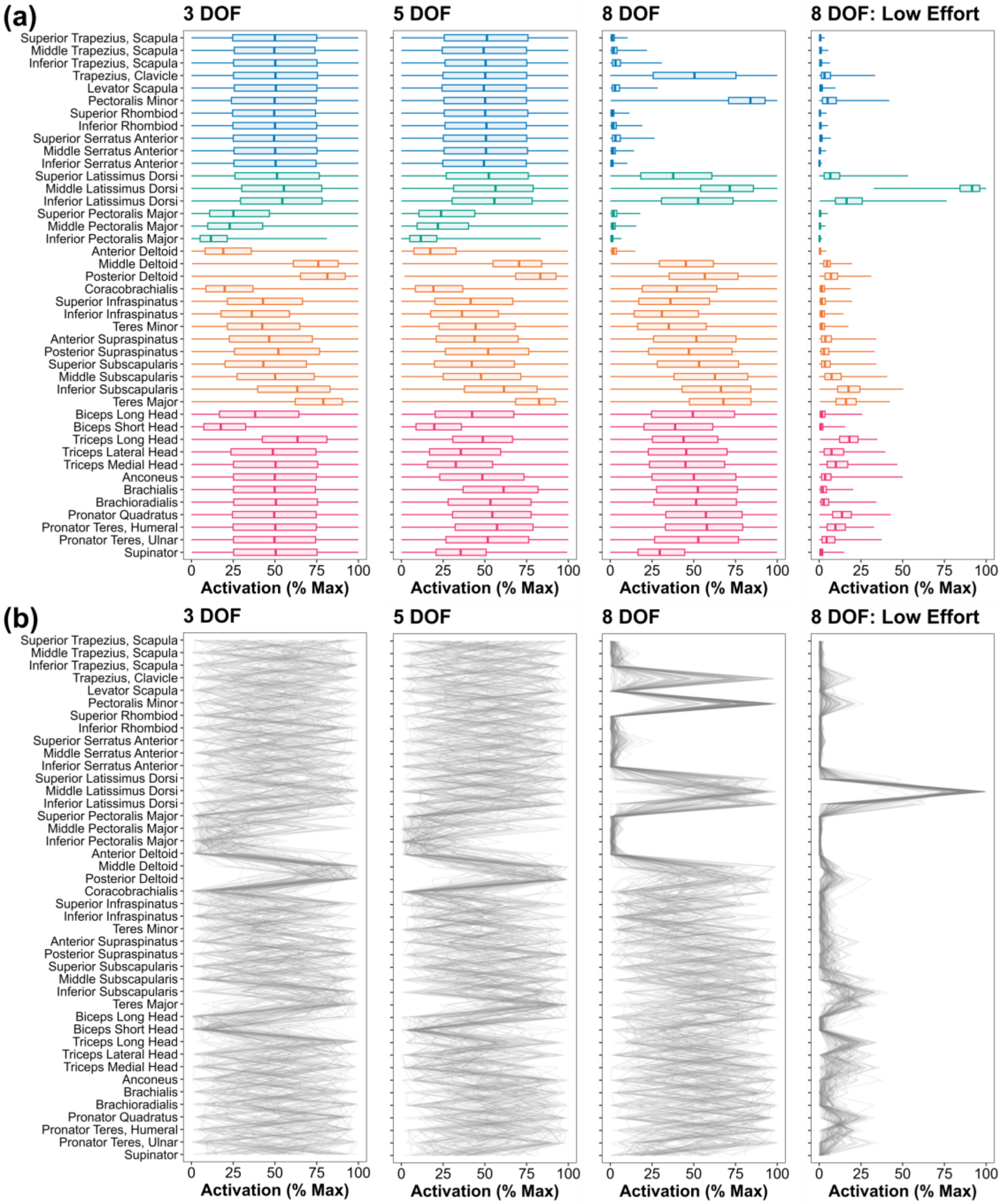


**Figure S4:** (a) Distribution of feasible solutions for a downwards 50 N exertion across models of different number of DOF with the reserve actuators set at 5 Nm for the SC and AC joints, but 1 Nm at all other DOF. Please note this is the same task as presented in Figure 2 of the main manuscript but using reduced SC and AC reserve actuator strength (5 Nm vs. 10 Nm). The colours correspond to different muscle groups (blue: thoracoscapular, green: thoracohumeral, orange: scapulohumeral, pink: elbow). (b) A random sample of 100 solutions from the feasible solution space from the corresponding distributions above visualized as a flipped parallel coordinates plot. Each grey line is a single solution drawn from top to bottom (i.e., activation combination across muscles).
